# eIF2B-capturing viral protein NSs suppresses the integrated stress response

**DOI:** 10.1101/2021.06.07.447466

**Authors:** Kazuhiro Kashiwagi, Yuichi Shichino, Tatsuya Osaki, Ayako Sakamoto, Madoka Nishimoto, Mari Takahashi, Mari Mito, Friedemann Weber, Yoshiho Ikeuchi, Shintaro Iwasaki, Takuhiro Ito

## Abstract

Various stressors such as viral infection lead to the suppression of cap-dependent translation and the activation of the integrated stress response (ISR), since the stress-induced phosphorylated eukaryotic translation initiation factor 2 [eIF2(αP)] tightly binds to eIF2B to prevent it from exchanging guanine nucleotides on unphosphorylated eIF2. Sandfly fever Sicilian virus (SFSV) evades this cap-dependent translation suppression through the interaction between its nonstructural protein NSs and host eIF2B. Our cryo-electron microscopy (cryo-EM) analysis revealed that SFSV NSs binds to the α-subunit of eIF2B in a competitive manner with eIF2(αP). Together with SFSV NSs, eIF2B exhibits normal nucleotide exchange activity even in the presence of eIF2(αP). A genome-wide ribosome profiling analysis clarified that SFSV NSs in human cultured cells attenuates the ISR. Furthermore, SFSV NSs exhibited neuroprotective effects against the ISR-inducing stress. Since the ISR inhibition is beneficial in various neurological disease models, SFSV NSs is promising as a therapeutic ISR inhibitor.

## INTRODUCTION

Upon various stresses, eukaryotic cells activate a common pathway called the integrated stress response (ISR), a homeostatic pathway necessary for organismal fitness. In vertebrates, four protein kinases, PKR, PERK, GCN2, and HRI, which are activated by different stress stimuli, commonly phosphorylate the α-subunit of eukaryotic initiation factor 2 (eIF2). Diverse stress signals are thus integrated into the phosphorylation of eIF2, and activate the common downstream pathway (Pakos-Zebrucka, et al., 2016).

eIF2 is a heterotrimeric G-protein that delivers an initiator methionyl-tRNA (Met-tRNA_i_) to the 40S ribosomes in a GTP-dependent manner, and is also involved in the start codon recognition process on the ribosomes. After the recognition, eIF2 is released from the ribosomes as the GDP-bound form, and regenerated by its specific guanine nucleotide exchange factor, eIF2B, a heterodecameric complex of two copies each of the α-, β-, γ-, δ- and ε-subunits (Merrick & Pavitt, 2018). However, phosphorylated eIF2 [eIF2(αP)] binds to eIF2B in an alternative nonproductive mode, and interferes with the guanine nucleotide exchange reaction of eIF2B (Kashiwagi et al., 2019; Kenner et al., 2019). Since eIF2B is less abundant than eIF2 in cells, the phosphorylation of a portion of eIF2 can sufficiently inhibit the catalytic activity of eIF2B (Merrick & Pavitt, 2018). This inhibition limits the supply of active GTP-bound eIF2, resulting in global attenuation of protein synthesis and selective translation of stress-related mRNAs (Wek, 2018).

Infection by various DNA and RNA viruses is one of the drivers of the ISR program. PKR is activated by double-stranded RNA derived from the virus, and phosphorylates eIF2 to shut off global protein synthesis (Stern-Ginossar et al., 2019). On the other hand, viruses have also evolved several strategies to evade this host defense mechanism. For example, poliovirus induces the degradation of PKR (Black et al., 2019), vaccinia virus blocks the phosphorylation of eIF2 by producing a pseudo-substrate for PKR (Kawagishi-Kobayashi et al., 1997; Sharp et al., 1997), and herpes simplex virus induces the dephosphorylation of eIF2(αP) (He et al., 1997). Recently, the nonstructural protein NSs of sandfly fever Sicilian virus (SFSV) was shown to exploit a novel mechanism to evade the protein synthesis arrest (Wuerth et al., 2020). SFSV belongs to the genus *Phlebovirus* (order *Bunyavirales*) and is endemic in the Mediterranean region (Depaquit et al., 2010). During the SFSV infection, PKR activation and eIF2 phosphorylation are both induced. However, the viral NSs protein counteracts the attenuation of cap-dependent translation. Since SFSV NSs binds to eIF2B, it may force eIF2B into the productive mode (Wuerth et al., 2020). Proteins from beluga whale coronavirus and Aichi picornavirus also reportedly target eIF2B (Rabouw et al., 2020). However, their molecular mechanisms have not been elucidated in detail.

The property of SFSV NSs, which directly binds to eIF2B and prevents the attenuation of cap-dependent translation, is somewhat similar to that of ISRIB, a small molecule that acts as an allosteric eIF2(αP) antagonist, binding eIF2B to disfavor its interaction with eIF2(αP) (Schoof et al., 2021; Zyryanova et al., 2021). Interestingly, ISRIB has shown promising effects in various animal models of neuropathological conditions, such as traumatic brain injury (Chou et al., 2017), amyotrophic lateral sclerosis (ALS) (Bugallo et al., 2020), Down syndrome (Zhu et al., 2019), and prion disease (Halliday et al., 2015), and even in the normal aging process (Krukowski et al., 2020).

## RESULTS AND DISCUSSISON

### SFSV NSs and eIF2(αP) bind to overlapping regions on eIF2B

We successfully purified the SFSV NSs protein and determined the complex structure of human eIF2B and SFSV NSs by cryo-electron microscopy (cryo-EM) at 2.4-Å resolution (Figures 1 and S1; Table 1). Two SFSV NSs molecules bind one eIF2B molecule, and are located between the α- and δ-subunits of eIF2B (Figure 1A). The interfaces for SFSV NSs partially overlap with those of the phosphorylated eIF2α subunit in the eIF2B•eIF2(αP) complex (Zyryanova et al., 2021), but there is no similarity between the phosphorylated eIF2α and SFSV NSs in terms of their three-dimensional structures and interaction modes with eIF2B (Figure 1B).

**Figure 1.**
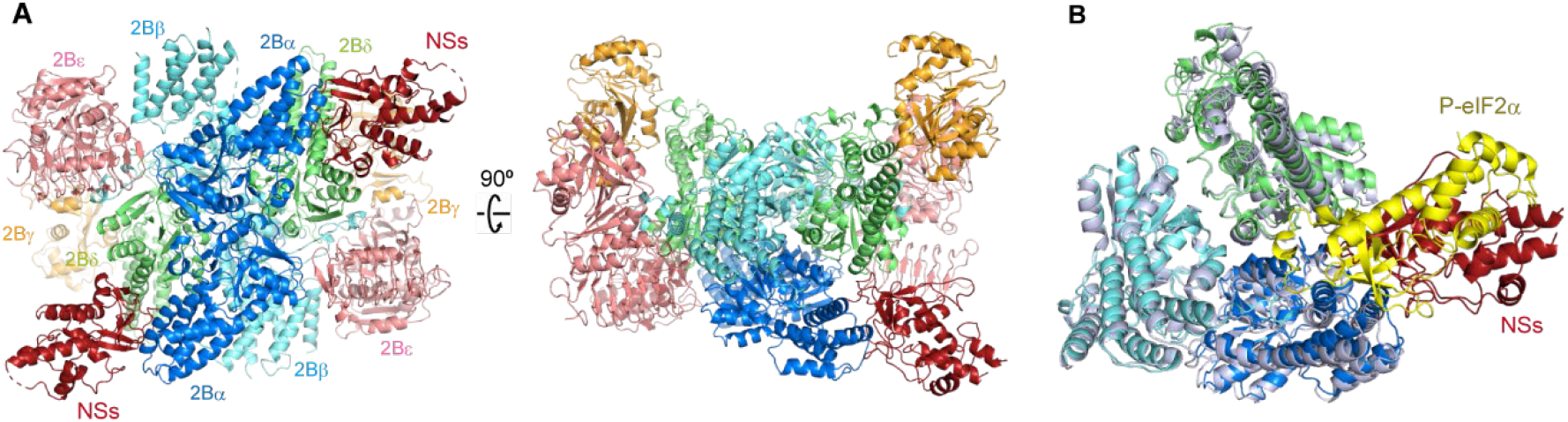
Cryo-EM structure of the human eIF2B•SFSV NSs complex. (A) Overall structure of the human eIF2B•SFSV NSs complex (eIF2Bα: blue; eIF2Bβ: cyan; eF2Bγ: orange; eIF2Bδ: lime; eIF2Bε: pink; SFSV NSs: maroon). (B) Overlay showing a comparison of the interactions through the α-δ groove of eIF2B in the eIF2B•SFSV NSs complex and in the eIF2B•eIF2(αP) complex (PDB: 7D44) (Zyryanova et al., 2021). The eIF2B subunits and the phosphorylated eIF2α subunit in the eIF2B•eIF2(αP) complex are shown in pale blue and yellow, respectively. Two structures are aligned with their eIF2Bβ subunit C-terminal domains. See also Figure S1.

**Table 1.** Cryo-EM data collection, refinement and validation statistics.

|  | eIF2B•SFSV NSs<br>(EMDB-xxxxx)<br>(PDB: XXXX) | eIF2B•SFSV NSs•1-eIF2<br>(EMDB-yyyyy)<br>(PDB: YYYY) | eIF2B•SFSV NSs•2-eIF2<br>(EMDB-zzzz)<br>(PDB: ZZZZ) |
| --- | --- | --- | --- |
| **Data collection and processing** |  |  |  |
| Magnification | 105,000 | 105,000 |  |
| Voltage (kV) | 300 | 300 |  |
| Electron exposure (e<sup>-</sup>/Å<sup>2</sup>) | 50 | 50 |  |
| Defocus range (μm) | -1.0 to -3.0 | -1.5 to -3.0 |  |
| Pixel size (Å) | 0.829 | 0.829 |  |
| Symmetry imposed | C1 | C1 |  |
| Initial particle images (no.) | 3,413,435 | 4,482,492 |  |
| Final particle images (no.) | 888,479 | 114,377 | 8,106 |
| Map resolution (Å) | 2.42 | 2.76 | 3.59 |
| FSC threshold | 0.143 | 0.143 | 0.143 |
| **Refinement** |  |  |  |
| Initial model used (PDB code) | 6O9Z | 6O9Z, 6O85 | 6O9Z, 6O85 |
| Model resolution (Å) | 2.7 | 2.9 | 3.7 |
| FSC threshold | 0.5 | 0.5 | 0.5 |
| Map sharpening *B* factor (Å<sup>2</sup>) | -65.3 | -56.1 | -56.4 |
| Model composition |  |  |  |
| Non-hydrogen atoms | 28736 | 33710 | 37714 |
| Protein residues | 3694 | 4545 | 5358 |
| *B* factors (Å<sup>2</sup>) |  |  |  |
| Protein | 77.30 | 101.76 | 152.71 |
| R.m.s. deviations |  |  |  |
| Bond lengths (Å) | 0.003 | 0.003 | 0.004 |
| Bond angles (°) | 0.588 | 0.583 | 0.610 |
| Validation |  |  |  |
| MolProbity score | 1.62 | 1.68 | 1.89 |
| Clashscore | 7.63 | 6.44 | 9.06 |
| Poor rotamers (%) | 1.53 | 0.03 | 0.09 |
| Ramachandran plot |  |  |  |
| Favored (%) | 97.69 | 95.29 | 94.01 |
| Allowed (%) | 2.31 | 4.67 | 5.99 |
| Disallowed (%) | 0.00 | 0.04 | 0.00 |
| CCmask | 0.82 | 0.88 | 0.84 |

2 See also Figure S1 and S2.

We were able to assign the amino (N)-terminal ~200 residues out of the 261 residues of SFSV NSs in the cryo-EM map (Figure S1D). The N-terminal region of SFSV NSs contains a β-sheet, followed by a helical core. The arrangement of the helices is similar to that in the core domain of the Rift Valley fever virus (RVFV) NSs protein (Barski et al., 2017), another virus in the genus *Phlebovirus*, that however targets PKR (Wuerth and Weber, 2016) (Figure S1E). The interaction of SFSV NSs with eIF2B is mediated by the N-terminal β-sheet region, and the β-sheets and protruding loops form a concave interface that embraces helix α3 in the N-terminal domain of the eIF2Bα subunit. The very N-terminus of SFSV NSs is also involved in the interaction with eIF2Bα (Figure S1F). Therefore, if a peptide artificially extends the N-terminus, it would interfere with the binding to eIF2Bα. This explains the previous observation that the N-terminally tagged version of the SFSV NSs protein completely lost its activity and the ability to interact with eIF2B (Wuerth et al., 2020). The eIF2B interaction with SFSV NSs is mostly mediated by the α-subunits, and does not induce a large movement in the eIF2B subunits as compared with the apo eIF2B structure (Zyryanova et al., 2021) (Figure S1G). This is in stark contrast to the phosphorylated eIF2α, which binds like a wedge between the α- and δ-subunits of eIF2B and interacts with both subunits extensively, resulting in the structural rearrangement of the eIF2B subunits (Schoof et al., 2021; Zyryanova et al., 2021) (Figures 1B and S1G).

One feature of the interface between eIF2B and SFSV NSs is the abundance of aromatic residues on the SFSV NSs side. These residues form two clusters (clusters 1 [c1] and 2 [c2]) and bury spaces between the helices of the eIF2Bα subunit (Figure 2A). To validate the importance of these aromatic residues, they were substituted with alanine and the dissociation constants with eIF2B were measured by microscale thermophoresis (MST). Wild-type SFSV NSs exhibited tight binding to eIF2B, with a dissociation constant of 16.9 ± 2.8 nM (Figure 2B). The alanine substitutions in c2 (SFSV NSs-c2-Ala-mut; Y79A and F80A) weakened the interaction by about six-fold, and the alanine substitutions in c1 (SFSV NSs-c1-Ala-mut; Y5A, F7A, and F33A) or in both clusters (SFSV NSs-c1+2-Ala-mut; Y5A, F7A, F33A, Y79A, and F80A) resulted in more than hundred-fold reductions of the interaction (Figure 2B).

**Figure 2.**
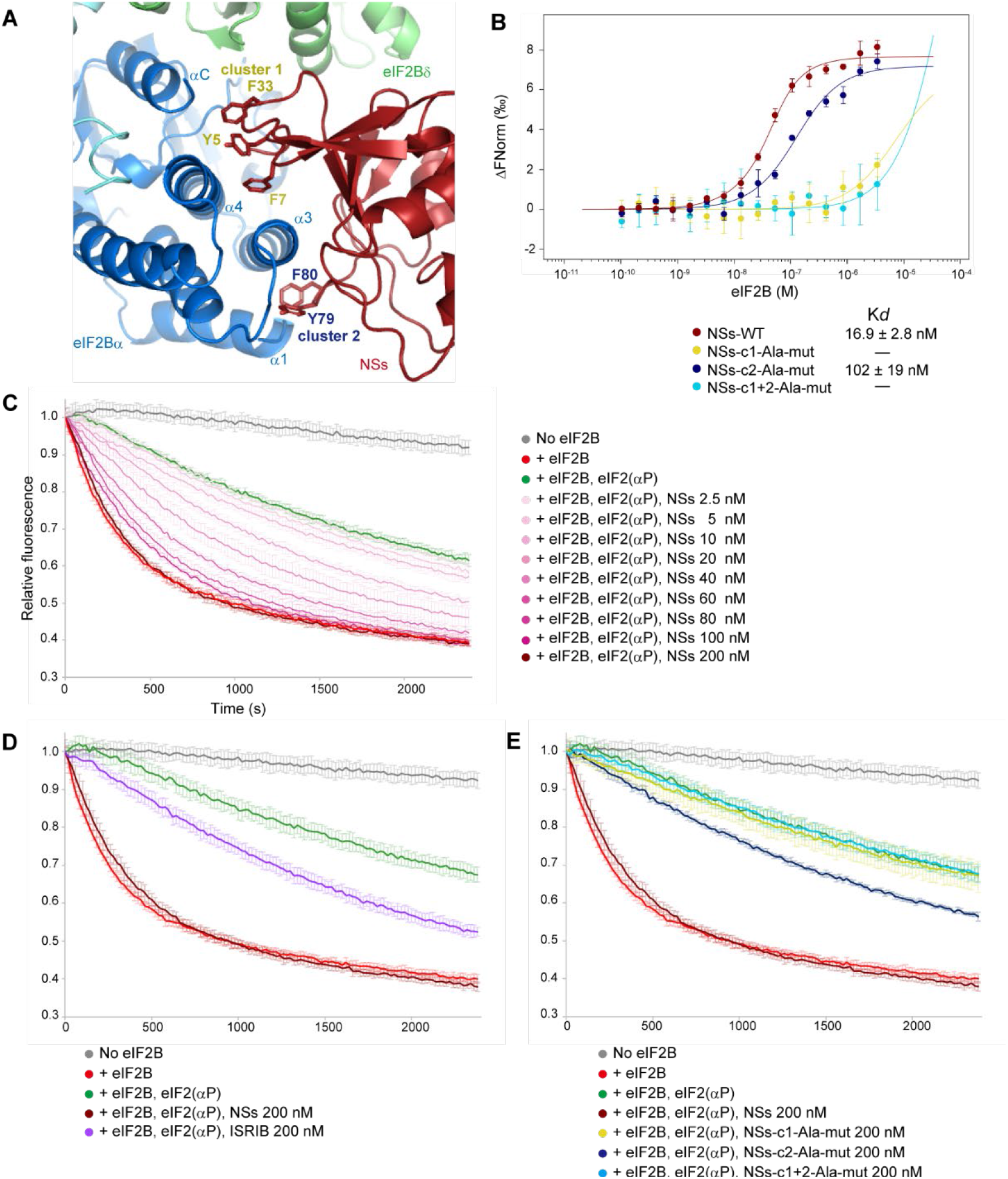
Aromatic clusters of SFSV NSs are important in the binding to eIF2B and the suppression of the inhibitory effect of eIF2(αP) (A) Two aromatic clusters of SFSV NSs at the interface with eIF2B. Aromatic cluster 1 (c1: Y5, F7, and F33) and aromatic cluster 2 (c2: Y79 and F80) grasp the α3 helix of the eIF2Bα subunit from both sides and bury the space between the helices of the eIF2Bα subunit. (B) MST analysis between eIF2B and SFSV NSs. Fluorescently-labeled SFSV NSs (100 nM) was mixed with an equal volume of a 16-step serial dilution of 6.8 μM eIF2B, and the microscale thermophoresis was measured. Plots of the wild type SFSV NSs (maroon), and the alanine-substituted mutants on cluster 1 (SFSV NSs-c1-Ala-mut: Y5A, F7A, and F33A; yellow), cluster 2 (SFSV NSs-c2-Ala-mut: Y79A and F80A; blue), and both clusters (SFSV NSs-c1+2-Ala-mut: Y5A, F7A, F33A, Y79A, and F80A; cyan) are shown. The error bars represent standard deviations at each concentration (n = 3). (C–E) Guanine nucleotide exchange assay. Non-phosphorylatable eIF2(αS51A) (final 150 nM) was loaded with BODIPY-GDP and fluorescent signals were read every 20 s. In both panels, the grey lines are the measurements without eIF2B, and other measurements were started by the addition of eIF2B (final 40 nM). The red and green lines are the measurements without and with eIF2(αP) (final 1.5 μM), respectively. In the experiments shown in panel (C), wild type SFSV NSs was included at various concentrations (0–200 nM) in the reaction solution containing fluorescent-eIF2(αS51A) and eIF2(αP). In the experiments shown in panels (D) and (E), ISRIB (D) or the mutants of SFSV NSs (c1-Ala-mut, c2-Ala-mut, and c1+2-Ala-mut) (e) were included into the reaction solutions at 200 nM. The same controls (No eIF2B, eIF2B, eIF2B+eIF2(αP), and eIF2B+eIF2(αP)+SFSV NSs at 200 nM) were used in (D) and (E). The error bars represent standard deviations at each time point [n = 3 for eIF2B+eIF2(αP)+SFSV NSs at 100 nM in (C), n = 4 for eIF2B+eIF2(αP)+SFSV NSs at 20, 40, 60 nM in (C), eIF2B+eIF2(αP), eIF2B+eIF2(αP)+SFSV NSs-c1+2-Ala-mut in (E), and n = 5 for the rest of the experiments].

### SFSV NSs binding suppresses the inhibitory effect of eIF2(αP) on eIF2B

To understand the effect of this binding on the eIF2(αP)-mediated eIF2B inhibition, we performed GDP exchange experiments. The results revealed that the guanine nucleotide exchange activity of eIF2B was inhibited in the presence of eIF2(αP), as previously described. The inclusion of SFSV NSs to the reaction resulted in the suppression of this inhibitory effect of eIF2(αP) in a concentration-dependent manner (Figure 2C). While the suppression by ISRIB was only partial, as previously reported^18^, that by SFSV NSs was more robust and almost nullified the inhibitory effect of eIF2(αP) (Figures 2C and 2D). This may reflect the difference in their eIF2B-binding modes, in which SFSV NSs physically covers the interface for eIF2(αP), but ISRIB does not. The suppression was weakened or canceled by the alanine substitutions of the aromatic clusters of SFSV NSs, and their effects correlated well with their binding affinities to eIF2B (Figure 2E). This indicated that (i) the suppression of the inhibitory effect of eIF2(αP) depends on the binding of SFSV NSs to eIF2B, and (ii) the binding excludes eIF2(αP) from eIF2B, thus protecting eIF2B from the inhibitory action of eIF2(αP). In addition, these results demonstrated that the suppression strength of SFSV NSs can be controlled as intended, by introducing appropriate mutations at the eIF2B-interacting residues.

The currently proposed mechanism is apparently inconsistent with the previous observation that SFSV NSs does not interfere with eIF2(αP) binding to eIF2B (Wuerth et al., 2020). Even though the reason for this discrepancy is not clear, some fraction of eIF2B may bind one SFSV NSs molecule and one eIF2(αP) molecule simultaneously, through two distant interfaces around two symmetrically positioned eIF2Bα subunits.

### Binding of SFSV NSs is compatible with the catalytic activity of eIF2B

We also analyzed the mixture of human eIF2B, unphosphorylated eIF2, and SFSV NSs, and obtained the cryo-EM structures of the eIF2B•SFSV NSs•unphosphorylated eIF2 ternary complex (Figure 3). Most of the ternary complex particles contained only one molecule of eIF2 (Figure 3A), but small fractions of those containing two eIF2 molecules were also observed (Table 1; Figure S2). There are no noticeable structural differences in eIF2B or SFSV NSs between these two ternary complexes. These structures revealed that SFSV NSs and the unphosphorylated eIF2 can bind to eIF2B simultaneously and there is no direct interaction between them (Figure 3B). The SFSV NSs binding does not seem to affect the unphosphorylated eIF2α binding, because the SFSV NSs-bound eIF2B is able to accommodate eIF2 without major conformational changes. The only exception is the quite small closure of the interface for unphosphorylated eIF2 (Figure S2D), as observed in the comparison of the eIF2B•unphosphorylated eIF2 complex structure with the eIF2B apo structure (Zyryanova et al., 2021). The conformation of eIF2 is essentially the same as that in the eIF2B•unphosphorylated eIF2 complex structures, and the nucleotide-binding eIF2γ subunit is trapped in the nucleotide-free state, where the switch 1 region is widely open (Kashiwagi et al., 2019; Kenner et al., 2019) (Figure S2E). Therefore, the binding of SFSV NSs does not seem to affect the catalytic activity or the substrate recognition process of eIF2B, and the binding of unphosphorylated eIF2 also does not seem to affect the binding of SFSV NSs.

**Figure 3.**
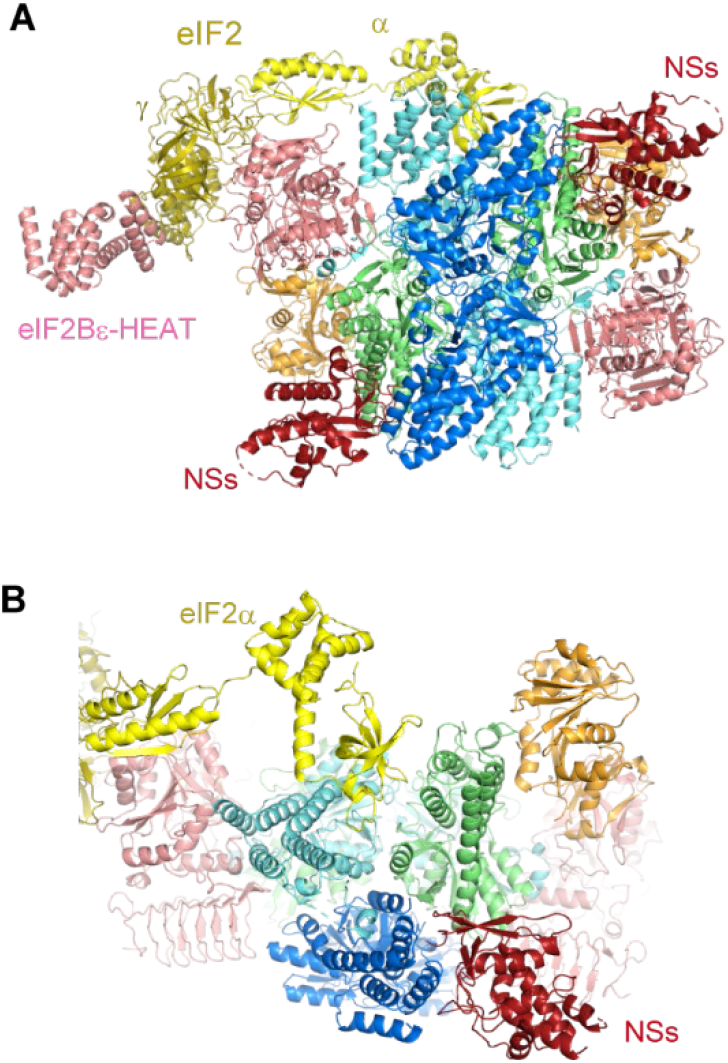
Cryo-EM structure of the human eIF2B•SFSV NSs•unphosphorylated eIF2 complex. (A) Overall structure of the human eIF2B•SFSV NSs•unphosphorylated eIF2 (one eIF2-bound) complex. eIF2B and SFSV NSs are color-coded as in Figure 1. The eIF2α and eIF2γ subunits are colored yellow and olive, respectively. (B) Close-up view of the interfaces for SFSV NSs and eIF2α. These two molecules can bind eIF2B simultaneously, and there is no direct interaction between them. See also Figure S2.

### SFSV NSs attenuates the ISR in cellular contexts

The inhibition of eIF2B by eIF2(αP) results in the global reduction of protein synthesis. To observe the cellular effect of SFSV NSs on overall translation during stress, human embryo kidney (HEK) 293 cells were treated with thapsigargin (Tg) to induce endoplasmic reticulum (ER) stress, activate PERK, and lead to eIF2α phosphorylation and consequently the ISR. The Tg treatment shut off global translation, as monitored by the metabolic labeling of newly synthesized proteins with *O*-propargyl-puromycin (OP-puro) (Iwasaki and Ingolia, 2017), in a dose-dependent manner (Figures. 4A and 4B). Strikingly, the ectopic expression of wild-type SFSV NSs (Figure S3A) rescued the translation repression induced by Tg, whereas, in stark contrast, the eIF2B-unbound mutant SFSV NSs (c1+2-Ala-mut) lost this activity (Figures 4A and 4B).

**Figure 4.**
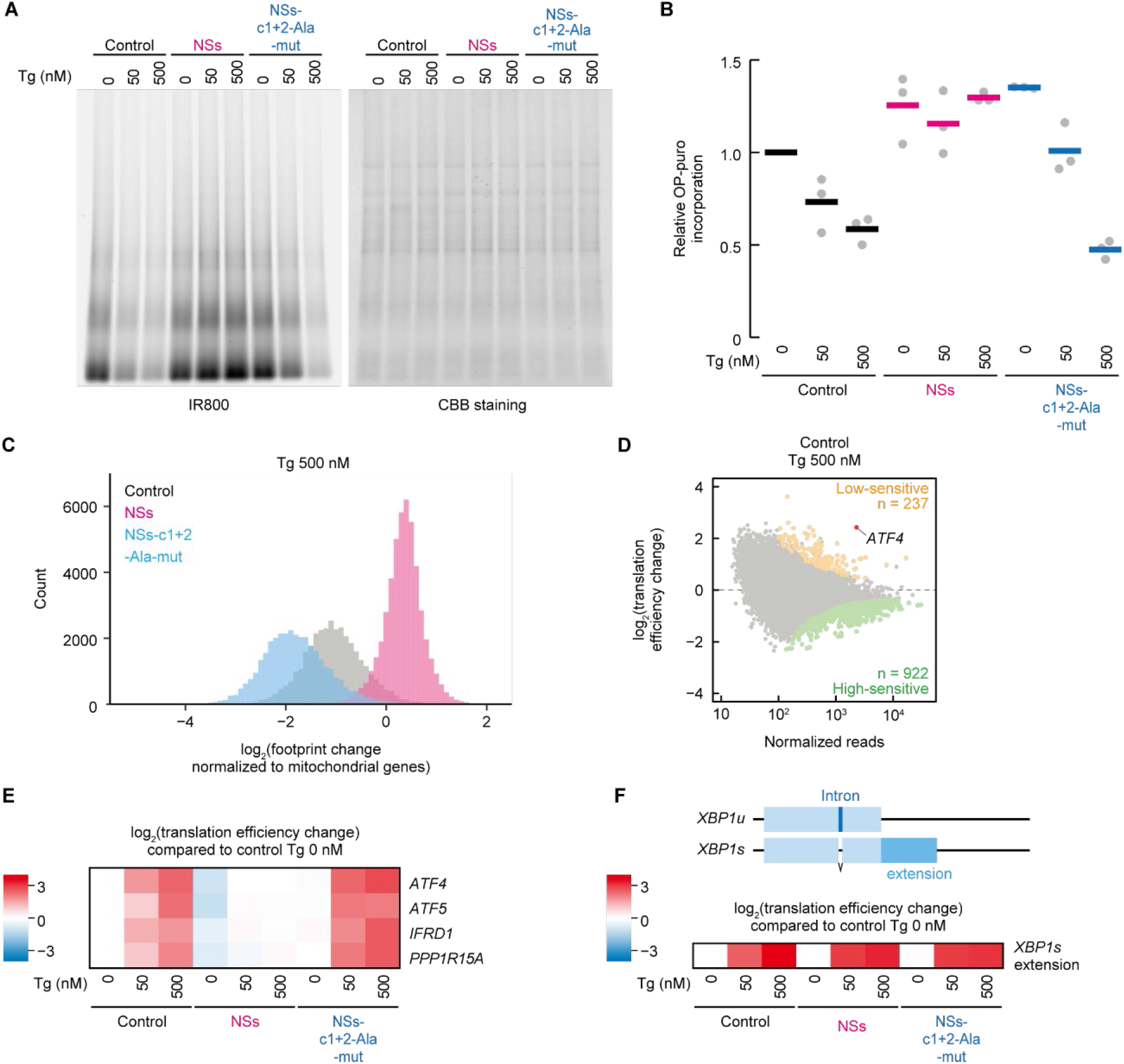
SFSV NSs suppresses translational impacts induced by thapsigargin. (A) Global protein synthesis rate measured by OP-puro labeling in control, SFSV NSs-expressing, or SFSV NSs-c1+2-Ala-mut-expressing cells. Cells were also treated with 50 or 500 nM Tg. Representative images of labeled nascent proteins (IR800 signal) and total protein with Coomassie Brilliant Blue (CBB) staining are shown. (B) Quantification of nascent proteins labeled with OP-puro, normalized by total protein in (A). Data of three replicates (points) and the mean (bar) are shown. (C) Histogram of the number of transcripts along the footprint change in cells treated with 500 nM Tg. Data were normalized to the mean of footprint change of mitochondrial genome-encoded genes (used as internal spike-ins). Bin width is 0.1. (D) MA (M, log ratio; A, mean average) plot of the translation efficiency change in control vector-transfected cells treated with 500 nM Tg. High-sensitive and low-sensitive mRNAs (defined as false discovery rate [FDR] < 0.05) are highlighted. (E and F) Heatmap of translation efficiency changes on *ATF4, ATF5, IFRD1, PPP1R15A* (E), and *XBP1s* extension (F), compared to control vector-transfected cells treated with 0 nM Tg. Schematic representation of the *XBP1* gene is shown on the top of (F). The intron in *XBP1u* mRNA is spliced to produce the C-terminally extended protein upon ER stress. See also Figures. S3 and S4.

The SFSV NSs-mediated recovery of translation repression in the ISR was also ensured in individual transcripts. To overview the landscape of translational output in the Tg-induced ISR and its recovery by SFSV NSs, we employed ribosome profiling (Ingolia et al., 2009). Congruent with the OP-puro results, we observed that the Tg-mediated translation repression across the transcriptome was recovered by wild-type SFSV NSs, but not by mutant SFSV NSs (Figures 4C, S3B, and S3C).

To further investigate the net alteration of translation over the transcriptome, which may be the ultimate consequence of ER stress (Pakos-Zebrucka et al., 2016), we evaluated the translational efficiency by the ribosome footprint abundance normalized by RNA sequencing (RNA-Seq). The translation efficiency changes in the ISR were not uniform among the transcripts, and a group of mRNAs, such as ribosomal protein mRNAs, were highly sensitive to Tg (Figures 4D and S3D). The translational recovery of these high-sensitivity mRNAs was accentuated upon wild-type SFSV NSs expression (Figure S3E).

Along with the suppression of protein synthesis, the phosphorylation of eIF2α simultaneously entails the translational activation of a subset of mRNAs to respond to the stress (Wek, 2018). This transcript-selective activation is associated with upstream ORFs (uORFs) in the 5’ UTR, which normally trap the scanning ribosome to block the complex reaching the main ORF further downstream. The reduced availability of the eIF2•GTP•Met-tRNA_i_ ternary complex evokes “leaky” scanning to skip the uORFs and then drives translation from the downstream main ORF. *ATF4*, a stress-inducible transcription factor, is a remarkable example of translational stimulation through uORFs (Pakos-Zebrucka et al., 2016). Indeed, upon Tg treatment, the translation from the main *ATF4* ORF was increased (Figures 4D, 4E, and S4A). In contrast, wild-type SFSV NSs expression did not activate *ATF4* translation, even with Tg treatment (Figure 4E). In addition to *ATF4*, uORF-bearing mRNAs, including *ATF5, IFRD1*, and *PPP1R15A*, were susceptible to similar translation activation in the ISR (Andreev et al., 2015) (Figure 4E). SFSV NSs also inhibited the activation of these mRNAs (Figure 4E). On the other hand, the eIF2B-unbound mutant SFSV NSs did not phenocopy the impact of wild-type SFSV NSs (Figures 4E and S4A).

In addition to PERK activation and subsequent eIF2α phosphorylation, ER stress also drives alternative branches of the stress response pathway, such as IRE1 activation and the subsequent *XBP1* mRNA splicing (Yoshida et al., 2001; Calfon et al., 2002), generating the *XBP1s* (spliced) mRNA with an extended ORF. The Tg-induced footprint accumulation in the extended ORF demonstrated the proper splicing of the mRNA and IRE1 activation even under the SFSV NSs expression (Figures 4F and S4B), and thus the high specificity of SFSV NSs in antagonizing the eIF2(αP) branch, but not the IRE1 branch of the ER stress response.

Taken together, our data demonstrate the clear correspondence among the cellular functions of SFSV NSs and its mutants to their structural and biochemical propensities.

### SFSV NSs protects rat hippocampal neurons and human iPS cell-derived motor neurons from ISR-inducible stress

Considering that ISRIB shows beneficial effects in various neuropathological models (Halliday et al., 2015; Chou et al., 2017; Zhu et al., 2019; Bugallo et al., 2020), it is tempting to speculate that SFSV NSs also effectively works in such conditions. To assess the neuroprotective activity of SFSV NSs under the ISR-inducible stress conditions, we prepared primary cultures of hippocampal neurons from rat embryos, and motor neurons differentiated from human induced pluripotent stem (iPS) cells (Osaki et al., 2020) (Figure 5). The Tg treatment of rat primary hippocampal neurons decreased the length of their neurites, as expected (Figures 5A and 5B). To quantify the arborization of neurites, a Sholl analysis was performed (Figure 5C). The expression of SFSV NSs significantly attenuated the neurite degradation triggered by Tg, whereas that of the mutant SFSV NSs (c1+2-Ala-mut) did not indicate neuroprotective activity (Figures 5B and 5D). Immunostaining of ATF4 in the control neurons demonstrated that the Tg treatment significantly upregulated its expression (Figure 5E). Consistent with the phenotypic effect, SFSV NSs significantly suppressed the induction of ATF4 expression, in contrast to the mutant SFSV NSs (Figure 5E). The effect of SFSV NSs was further confirmed with human iPS cell-derived motor neurons (Figures 5F–5J). SFSV NSs efficiently suppressed the induction of ATF4 even with the Tg treatment, but the mutant SFSV NSs did not (Figure 5I). In addition, SFSV NSs promoted the axon lengthening and increased the density of elongated axons characterized by the Sholl analysis, as compared to the control and mutant SFSV NSs (Figure 5J). These results showed that SFSV NSs can also suppress the ISR in neurons during the ISR-inducible stress, thus providing the neuroprotective effect.

**Figure 5.**
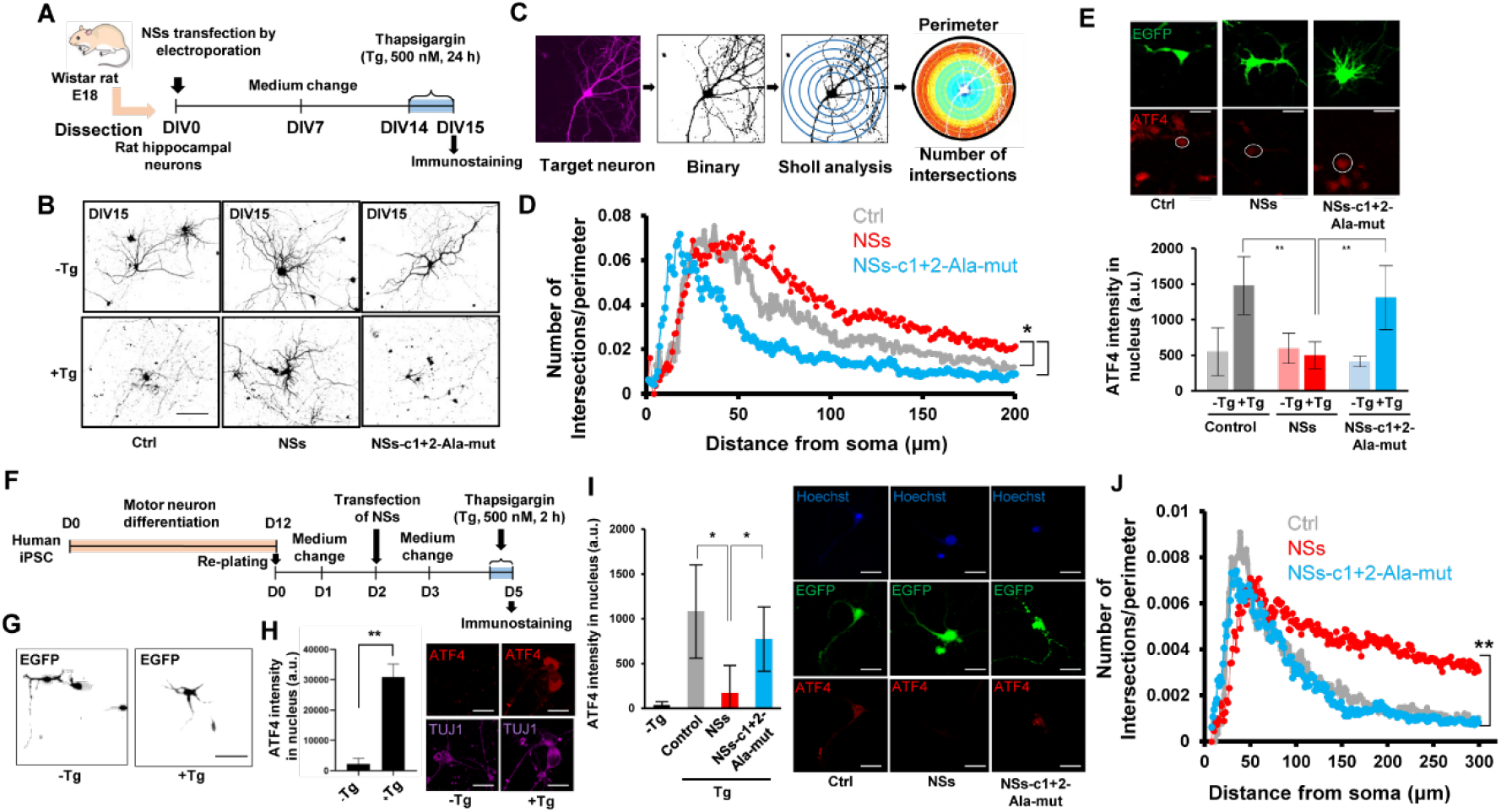
SFSV NSs in rat primary hippocampal neurons or human iPS-derived motor neurons attenuates the ISR induction and exhibits neuroprotective effects. (A) Schematic illustration of thapsigargin treatment (500 nM) for rat primary hippocampal neurons. The neurons were transfected with the control plasmid, SFSV NSs-expression plasmid, and SFSV NSs-c1+2-Ala-mut-expression plasmid by electroporation. (B) Representative images of hippocampal neurons with or without Tg treatment after 24 h. Tg treatment shortened the neurites in the control and SFSV NSs-c1+2-Ala-mut expressing cells, whereas SFSV NSs counteracted Tg. Scale bar = 100 μm. (C) Schematic representation of the Sholl analysis. (D) The Sholl analysis revealed that SFSV NSs protected the arborization of neurons. n = 5. (E) SFSV NSs downregulated ATF4 expression induced by Tg (500 nM, 3h), as compared to the control and SFSV NSs-c1 +2-Ala-mut. n = 7. *, *p* < 0.05. Scale bar = 10 μm. (F) Motor neuron differentiation from human iPS cells under the conditions of SFSV NSs transfection and drug administration. After motor neurons were replated onto glass cover slips, they were transfected with the control plasmid, SFSV NSs-expression plasmid, and SFSV NSs-c1+2-Ala-mut-expression plasmid by Lipofectamine 3000 for 2 days. After an additional 3 days, the neurons were treated with thapsigargin (Tg; 500 nM) for 2 h, followed by fixation and immunostaining of ATF4. (G) Representative images of motor neurons before and after Tg treatment. Scale bar = 50 μm. (H) Tg significantly induced the stress response, as represented by ATF4 upregulation in motor neurons. n = 3. Scale bar = 10 μm. **, *p* < 0.01. (I) ATF4 intensity in control, SFSV NSs-expressing, and SFSV NSs-c1+2-Ala-mut-expressing motor neurons after Tg treatment. In SFSV NSs-expressing motor neurons, ATF4 was significantly downregulated, whereas mutant SFSV NSs expression did not alter ATF4 protein synthesis as compared to the control. n = 3. *, *p* < 0.05. Scale bar = 10 μm. (J) Sholl analysis in motor neurons. SFSV NSs expression increased the length of axons as compared to the control and SFSV NSs-c1+2-Ala-mut expression. n = 5. *, *p* < 0.05. **, *p* < 0.01. (H) Student’s t test; (D, I, J) One-way ANOVA; (E) Two-way ANOVA. Error bars ± SD.

### Concluding remarks

We have solved the cryo-EM structures of the eIF2B•SFSV NSs complex and the eIF2B•SFSV NSs•unphosphorylated eIF2 ternary complex, and analyzed the catalytic effect of SFSV NSs binding to eIF2B upon its inhibition by eIF2(αP). The binding of SFSV NSs sterically blocks the inhibitory interaction between eIF2B and eIF2(αP), whereas it does not affect the catalytic interaction between eIF2B and unphosphorylated eIF2 (Figures 1 and 3). As a result, eIF2B becomes insensitive to the inhibitory effect of eIF2(αP) and continues to catalyze nucleotide exchange on residual unphosphorylated eIF2, even in the presence of high levels of eIF2(αP) (Figure 2). This is likely to be the molecular mechanism by which SFSV enables viral protein synthesis against the PKR-derived host cell defense.

The suppression of the ISR by SFSV NSs was observed in several kinds of animal cells: HEK293 cells, rat hippocampal neurons, and human iPS-derived motor neurons (Figures 4 and 5). Interestingly, the supplementation of SFSV NSs to neural cells exhibited neuroprotective effects under the ISR-inducible stressed conditions (Figure 5). The currently identified direct block mechanism of SFSV NSs against eIF2(αP) on eIF2B was more effective than the allosteric antagonism by ISRIB (Schoof et al., 2021; Zyryanova et al., 2021) (Figure 2D). Therefore, these observations raise the possibility that, like ISRIB, SFSV NSs could also show beneficial effects in neuropathological conditions. Future studies on the *in vivo* effect of SFSV NSs, administered as a drug in the respective animal models, may reveal the potential therapeutic utility of this strong ISR inhibitor.

## Acknowledgements

We thank David Ron and Alisa F. Zyryanova for providing detailed protocols for the guanine nucleotide exchange experiments. We are grateful to all of the members of the Ito, Iwasaki, and Ikeuchi laboratories for constructive discussions, technical help, and critical reading of the manuscript.

T.I. was supported by Grants-in-Aid for Scientific Research (B) (JP19H03172) from the Japan Society for the Promotion of Science (JSPS), AMED-CREST (JP21gm1410001) from the Japan Agency for Medical Research and Development (AMED), and the BDR Structural Cell Biology Project, the Pioneering Projects (“Dynamic Structural Biology” and “Biology of Intracellular Environments”), and the All RIKEN Research Project (“Integrated life science research to challenge super aging society”) from RIKEN.

S. I. was supported by a Grant-in-Aid for Transformative Research Areas (B) “Parametric Translation” (JP20H05784) from the Ministry of Education, Culture, Sports, Science and Technology (MEXT), a Grant-in-Aid for Young Scientists (A) (JP17H04998), and a Grant-in-Aid for Challenging Research (Exploratory) (JP19K22406) from JSPS, an AMED-CREST (JP21gm1410001) grant from AMED, the Pioneering project (“Biology of Intracellular Environments”) and the All RIKEN Research Project (“Integrated life science research to challenge super aging society”) from RIKEN, and the Takeda Science Foundation.

Y.I. was supported by a Grant-in-Aid for Transformative Research Areas (B) “Parametric Translation” (JP20H05786) from MEXT, a Challenging Research (Pioneering) (JP20K20643) grant from JSPS, an AMED-CREST (JP21gm1410001) grant from AMED, and the Institute for AI and Beyond.

K.K. was supported by a Grant-in-Aid for Early-Career Scientists (JP18K14644) from JSPS. Y.S. was supported by a Grant-in-Aid for Early-Career Scientists (JP21K15023) from JSPS and by the Special Postdoctoral Researchers Program and Incentive Research Projects from RIKEN.

T. O. was supported by the Japan Society for the Promotion of Science (JSPS) with a Grant-in-Aid for Early-Career Scientists (20K20178), an AMED-P-CREATE (G02-53) grant, and the Takeda Science Foundation.

F.W. was supported by the Deutsche Forschungsgemeinschaft (DFG; German Research Foundation; project number 197785619–SFB 1021) and the Swedish Research Council (VR; no. 2018-05766).

DNA libraries were sequenced by the Vincent J. Coates Genomics Sequencing Laboratory at UC Berkeley, supported by an NIH S10 OD018174 Instrumentation Grant. This study was supported by the Support Unit for Bio-Material Analysis, RIKEN CBS Research Resources Division for Sanger sequencing and by the supercomputer HOKUSAI SailingShip in RIKEN ACCC for computations. Structural analysis was supported by the Platform Project for Supporting Drug Discovery and Life Science Research (Basis for Supporting Innovative Drug Discovery and Life Science Research [BINDS], JP21am0101082) from AMED.

## Author Contributions

K.K. and T.I. designed and performed cryo-EM and biochemical experiments. Y.S., M.M., and S.I. designed and performed ribosomal profiling experiments. T.O. and Y.I. designed and performed experiments in rat primary hippocampal neurons and iPS cell-derived motor neurons. F.W. constructed the expression vectors. A.S., M.N., and M.T. prepared the samples. The manuscript was written by K.K., Y.S., T.O., F.W., S.I., Y.I., and T.I., and all authors contributed to manuscript editing.

## RESOURCE AVAILABILITY

### Lead Contact

Further information and requests for resources and reagents should be directed to and will be fulfilled by the Lead Contact, Takuhiro Ito.

### Materials Availability

Plasmids generated in this study are available upon request to the Lead Contact.

### Data and Code Availability

The cryo-EM maps and the coordinates of the refined models have been deposited in the Electron Microscopy Data Bank (EMDB) and the Protein Data Bank (PDB) under the following accession numbers: eIF2B•SFSV NSs (EMDB: EMD-xxxxx, PDB: XXXX), eIF2B•SFSV NSs•1-eIF2 (EMDB: EMD-yyyyy, PDB: YYYY), eIF2B•SFSV NSs•2-eIF2 (EMDB: EMD-zzzzz, PDB: ZZZZ). The ribosome profiling and RNA-Seq (GSE174764) results from this study have been deposited in the National Center for Biotechnology Information (NCBI). All other data are available in the manuscript or supplementary materials. Materials are available from Y.I., S.I. and T.I. upon request.

### Method Details

#### Protein preparation

Human eIF2B and eIF2 were recombinantly expressed and purified as previously described (Kashiwagi et al, 2019; Zyryanova et al., 2021).

The SFSV NSs gene was cloned into the pET-28a vector (Novagen) and expressed with a C-terminal His-tag in the T7 Express competent *E. coli* strain (NEB). The cells were grown at 37 °C in LB medium supplemented with 0.2% (w/v) glucose. After the addition of 0.3 mM isopropyl-β-D-thiogalactopyranoside (IPTG) at A_600_ = 0.5, the cells were further grown overnight at 18 °C. The harvested cells were lysed in lysis buffer [20 mM HEPES-KOH buffer (pH 7.5), containing 150 mM KCl, 5% (v/v) glycerol, and 1 mM DTT] supplemented with protease inhibitors. After centrifugation, the supernatant was purified with cOmplete His-Tag Purification Resin (Roche) and by chromatography on a HiTrap Q HP column (GE Healthcare) with a 150–500 mM KCl gradient. The sample was further purified by chromatography on a Superdex 200 column (GE Healthcare) equilibrated with lysis buffer. The SFSV NSs mutants were also purified in the same manner.

#### Cryo-EM data collection and image processing

For the cryo-EM analyses, eIF2B and SFSV NSs were mixed at a molar ratio of 1:4, diluted to 60 nM for eIF2B and 240 nM for SFSV NSs, and supplemented with 0.06% (w/v) digitonin. For the eIF2B•SFSV NSs•unphosphorylated eIF2 ternary complex, eIF2B, SFSV NSs and eIF2 were mixed at a molar ratio of 1:3:4, diluted to 50 nM for eIF2B, 150 nM for SFSV NSs and 200 nM for eIF2, and supplemented with 0.06% (w/v) digitonin. The grids were prepared using a Vitrobot Mark IV (FEI) at 4 °C and 100% humidity. The samples (3 μl) were loaded onto Quantifoil R1.2/1.3 300 mesh copper grids (Quantifoil) covered with an in-house prepared amorphous carbon layer, incubated for 30 s, blotted for 3 s, and plunged into liquid ethane.

The datasets were collected with a Krios G4 transmission electron microscope (FEI) operated at 300 kV, using a K3 direct electron detector (Gatan) operated in the CDS-counting mode (0.829 Å/pixel), running at the RIKEN Center for Biosystems Dynamics Research in Yokohama, Japan. The total numbers of collected images were 12,341 for the eIF2B•SFSV NSs complex sample, and 13,858 for the eIF2B•SFSV NSs•unphosphorylated eIF2 complex sample. The collected images were fractionated to 50 frames, with a total dose of ~50 e^-^/Å^2^. Processing of cryo-EM data was performed with RELION-3.1 (Zivanov et al., 2020). The movie frames (from 2nd frame to 50th frame) were aligned with MotionCor2 (RELION’s own implementation) and the CTF parameters were estimated with CTFFIND-4.1 (Rohou and Grigorieff, 2015). Particles were automatically picked using template-free Laplacian-of-Gaussian (LoG) filters (diameter 150–300 Å).

For the eIF2B•SFSV NSs complex sample, the 3,413,435 automatically picked particles were extracted with two-fold binning (1.658 Å/pixel). After splitting the particles into three subsets, 2D classification was performed twice. For 3D classification, good classes from three subsets were merged and split again into three subsets, and a low-pass filtered (40 Å) map calculated from the crystal structure of *Schizosaccharomyces pombe* eIF2B (PDB: 5B04) (Kashiwagi et al., 2016) was used as the reference map. After the classification, the particles in the good classes from the three subsets that contained the SFSV NSs-like protrusion from the density of eIF2B (899,315 particles) were joined and re-extracted without rescaling (0.829 Å/pixel), and performed the 3D refinement, Bayesian polishing, and CTF refinement, and subjected to 3D refinement again. The refined particles were subjected to 3D classification again and the junk particles were discarded. The final number of particles is 888,479.

For the eIF2B•SFSV NSs•unphosphorylated eIF2 complex sample, the 4,482,492 automatically picked particles were similarly extracted, split into subsets, and subjected to 2D classification. For 3D classification, a low-pass filtered (40 Å) map of eIF2B•SFSV NSs was used as the reference map. The classes containing eIF2B, SFSV NSs, and additional density (615,804 particles) were merged into one dataset and classified again. The classes containing the eIF2-like density (125,080 particles) were re-extracted without rescaling, and performed the 3D refinement, Bayesian polishing, and CTF refinement, and subjected to 3D refinement again. The refined particles were subjected to 3D classification again and separated into the class containing one eIF2 molecule (114,137 particles) and the class containing two eIF2 molecules (8,106 particles).

To build a model of the eIF2B•SFSV NSs complex, the human eIF2B moiety in the cryo-EM structure of the eIF2B•phosphorylated eIF2α complex at 3.0-Å resolution (PDB: 6O9Z) (Kenner et al., 2019) was used as the initial model. At first, the model of eIF2B was manually fitted into the maps using UCSF Chimera (Pettersen et al., 2004). Map sharpening and model refinement were performed in PHENIX (Adams et al., 2010), and the models were further refined manually with Coot (Emsley et al., 2010). The model for SFSV NSs was built automatically with the Buccaneer software (Cowtan, 2006) in the CCP-EM suite (Burnley et al., 2017), and manually with Coot. For the eIF2B•SFSV NSs•unphosphorylated eIF2 complex, the eIF2B•SFSV NSs complex structure, eIF2 and the HEAT domain of the eIF2Bε subunit from the cryo-EM structure of the eIF2B•unphosphorylated eIF2 complex at 3.0-Å resolution (PDB: 6O85) (Kenner et al., 2019) were manually fitted into the maps with USCF Chimera, and refined with PHENIX and Coot.

The statistics for image processing and refinement are summarized in Table 1.

#### Microscale thermophoresis

Purified eIF2B and SFSV NSs were dialyzed against measurement buffer [20 mM HEPES-KOH buffer (pH 7.4), containing 150 mM KCl and 5 mM MgCl_2_] overnight and supplemented with 0.05% (v/v) Tween-20 after dialysis. The fluorescently-labeled SFSV NSs protein was prepared by mixing equal volumes of 200 nM SFSV NSs and 150 nM HIS Lite OG488-Tris NTA-Ni Complex (AAT Bioquest), and eIF2B was diluted to 6.8 μM. Microscale thermophoresis was measured at 23 °C with a Monolith NT.115 (Nanotemper), according to the manufacturer’s protocol. Each experiment was repeated three times for the dissociation constant calculations.

#### Guanine nucleotide exchange assay

The eIF2B guanine nucleotide exchange activity was measured as previously described (Zyryanova et al., 2021), with some modifications. Purified eIF2(αS51A) was loaded with BODIPY-FL-GDP (Invitrogen) in loading buffer [20 mM HEPES-KOH buffer (pH 7.4) containing 150 mM KCl, 1 mM DTT, 0.05 mg/ml bovine serum albumin (BSA), and 0.01% (v/v) Triton X-100] at room temperature. After supplementation with 2 mM MgCl_2_, the unbound fluorescent dye was removed and the buffer was exchanged to assay buffer [20 mM HEPES-KOH buffer (pH 7.4) containing 150 mM KCl, 2 mM MgCl_2_, 1 mM DTT, 0.05 mg/ml BSA, 0.01% (v/v) Triton X-100, and 1.5 mM GDP] using ProbeQuant G-50 Micro Columns (Cytiva). eIF2(αP) was prepared using PKR as previously described (Kashiwagi et al., 2019), and buffer exchange was performed as above. The proteins were premixed in a 384-well Low Volume Black Round Bottom Polystyrene NBS Microplate (Corning) at final concentrations of 150 nM fluorescently-labeled eIF2(αS51A), 0 or 1.5 μM eIF2(αP), 0–200 nM SFSV NSs, and 0 or 200 nM ISRIB (Sigma-Aldrich), and reactions were started by the addition of 40 nM eIF2B or assay buffer into the mixture at 25 °C. The fluorescence signal was read every 20 s in an EnVision 2104 plate reader (PerkinElmer), with excitation at 485 nm and emission at 535 nm, and each experiment was repeated three to five times.

#### Plasmid construction

##### pI.18-3xFLAG and SFSVNSs-c1+2-Ala-mut-3xFLAG

To construct pI.18-3xFLAG and pI.18-SFSVNSs-c1+2-Ala-mut-3xFLAG, the SFSV NSs gene in pI.18-SFSVNSs-3xFLAG (Wuerth et al, 2020) was removed or replaced with a PCR-amplified DNA fragment encoding the mutated SFSV NSs gene, respectively.

#### Nascent peptide labeling by OP-puro

HEK293 T-REx cells (Thermo Fisher Scientific) were cultured in a 24-well plate with DMEM + GlutaMAX-I (Thermo Fisher Scientific, 10566016) supplemented with 10% FBS (Sigma, F7524) at 5% CO_2_ and 37 °C, and transfected with 0.3 μg of pI.18-3xFLAG, pI.18-SFSVNSs-3xFLAG, or pI.18-SFSVNSs-c1+2-Ala-mut-3xFLAG plasmids, using the TransIT-293 Reagent (Mirus, MIR2704). After 48 h incubation, the cells were treated with thapsigargin (Nacalai Tesque) for 30 min at 37 °C. OP-puro labeling was performed as described previously (Iwasaki et al, 2019).

#### Ribosome profiling and RNA-Seq

##### Library preparation

HEK293 T-REx cells were cultured in 10-cm dishes with DMEM + GlutaMAX-I supplemented with 10% FBS at 5% CO_2_ and 37 °C, and transfected with 6 μg of pI.18-3xFLAG, pI.18-SFSVNSs-3xFLAG, or pI.18-SFSVNSs-c1+2-Ala-mut-3xFLAG plasmids, using the TransIT-293 Reagent. After 48 h incubation, the cells were treated with thapsigargin for 30 min at 37 °C. The ribosome profiling library was prepared according to the protocol described previously (McGlincy and Ingolia, 2017; Mito et al., 2020). Ribosome-protected RNA fragments ranging from 17–34 nt were excised from the gel after PAGE.

For RNA-Seq, RNA was extracted from the same lysate used for ribosome profiling with Trizol LS (Thermo Fisher Scientific, 10296-010) and a Direct-zol RNA microprep kit (Zymo Research, R2060), and 0.5 μg of the RNA was used for RNA-Seq library preparation with a TruSeq Stranded mRNA Library Prep Kit (Illumina). The libraries were sequenced on a HiSeq 4000 System (Illumina).

##### Data analysis

Sequence data were processed as previously described (McGlincy and Ingolia, 2017) with the following modifications. Read quality filtering and adapter trimming were performed with Fastp (Chen et al., 2018). After removing the non-coding RNA-mapped reads, the remaining reads were aligned to the human genome hg38 and assigned to the GENCODE Human release 32 reference, using STAR 2.7.0a (Dobin et al., 2013). For ribosome profiling, the offsets of the A site were determined to be 15 for 20–22 and 25–30 nt. For RNA-Seq, an offset of 15 was used for all mRNA fragments. Reads corresponding to the first and last five codons in CDS were excluded from the calculation.

Ribosome footprint changes were calculated with the DESeq2 package (Love et al., 2014). The values were renormalized to the mean of the footprint change of mitochondrial genome-encoded genes, which are used as the internal spike-in (Iwasaki et al., 2016). Translation efficiencies, which are ribosome profiling counts normalized by RNA-seq counts, were also measured by DESeq2. The significance was calculated by a likelihood ratio test in a generalized linear model. The gene ontology analysis was performed with iPAGE (Goodarzi et al., 2009). All custom scripts used in this study are available upon request.

#### SFSV NSs in rat primary hippocampal neurons and iPS-derived motor neurons

The hippocampal neurons were dissected from an E18 Slc:Wistar rat by using the standard dissection and dissociation protocol with 0.25% trypsin and DNase. Before plating, the cells were electroporated with pI.18-3xFLAG (control), pI.18-SFSVNSs-3xFLAG, or pI.18-SFSVNSs-c1+2-Ala-mut-3xFLAG and pCAG-EGFP (NEPA21, Nepa Gene). The hippocampal neurons were then plated on PLL-coated glass cover slips in a 24-well plate, at a density of 100,000 cells/well. The cells were maintained in Neurobasal medium (Thermo Fisher Scientific) supplemented with 2% B27 supplement (Gibco), 1% GlutaMAX, and 1% penicillin/streptomycin). After 2 weeks, thapsigargin (Tg, 500 nM) was added to the culture medium for either 3 or 24 h. The motor neuron differentiation protocol has been described previously (Osaki et al., 2020). Briefly, human induced pluripotent stem cells (409B2 from the RIKEN Cell Bank) were seeded in Matrigel-coated plates at 70–80% confluency, in mTeSR Plus medium (STEMCELL Technology) with 10 μM of Y-23632 (Rock inhibitor, Wako). After the cells reached 95% confluency, the neural differentiation was initiated in DMEM/F12 (Sigma) supplemented with 15% knockout serum replacement (KSR, Gibco), 1% GlutaMAX (Gibco), 1% non-essential amino acids (NEAA, Sigma), 10 μM SB431542 (Wako), and 100 nM LDN-193189 (Wako) for the initial 6 days of culture. The cells were treated with 1 μM retinoic acid (RA), 1 μM Smoothened agonist (SAG, Wako), 10 μM SU-5402 (Sigma), and 10 μM DAPT (Sigma) from the 4th to 12th days of culture. The motor neurons were then dissociated by Accutase (Innovative Cell Technology) for 20–25 min and replated in a Matrigel-coated 24-well plate containing glass cover slips, at a density of 100,000 cells/well. The motor neurons were maintained in Neurobasal medium containing 2% B27 supplement, 1% GlutaMAX, 1% penicillin/streptomycin, and brain-derived neurotrophic factor (BDNF, 20 ng/ml). At 2 days after the cells were replated, the neurons were transfected with pI.18-3xFLAG (control), pI.18-SFSVNSs-3xFLAG, or pI.18-SFSVNSs-c1+2-Ala-mut-3xFLAG together with the pCAG-EGFP plasmid, using Lipofectamine 3000. At 3 days after the transfection, thapsigargin (Tg, 500 nM) was added to the culture medium for 2 h. The neurons were fixed by 4% paraformaldehyde for 15 min and permeabilized by 0.1% Triton X-100 for 5 min, followed by blocking with 1% BSA. The cells were then incubated with an anti-ATF4 rabbit antibody (Cell Signaling Technologies), an anti-GFP mouse antibody (DHSB, 1:50) and an anti-beta 3 Tubulin mouse antibody (BioLegend, 1:1,000) in PBS at 4 °C overnight. On the next day, the cells were further incubated with Alexa Fluor 555-conjugated anti-rabbit IgG (Thermo Fisher Scientific, 1:1,000) and Alexa Fluor 647-conjugated anti-mouse IgG (Thermo Fisher Scientific, 1:1,000) in PBS at room temperature for 3 h with Hoechst 33342 for nuclear staining, followed by three washes with PBS. The fluorescent images were acquired with an Axio Observer inverted microscope (Zeiss) and a Nikon confocal microscope (A1R).

#### ATF4 intensity quantification

The rat primary neurons and motor neurons co-expressing SFSV NSs and GFP (target neurons) were chosen for the ATF4 expression analysis in the confocal and fluorescent images. The images of ATF4 channel (Red channel) containing target neurons were converted to 16-bit monochrome images, and then the ROI was determined according to the area of the nucleus indicated by Hoechst expression. The mean intensities of ATF4-staining signals in the ROIs were then calculated. All post-image analyses were conducted by using Fiji (http://imagej.nih.gov/ij/).

#### Sholl analysis

The rat primary hippocampal neurons and motor neurons expressing GFP were examined by a Sholl analysis. The images of neurons were binarized and clarified by removing noise (pixel < 2). After the center of cells (soma) was manually set, the Sholl analysis was performed by using the Sholl analysis plugin in Fiji. The macro command for the Sholl analysis is provided upon request.

#### Statistics

All of the details for the statistical tests with ‘n’ values are indicated in the relevant figure legends and method sections.

## Supplemental Information

**Figure S1.**
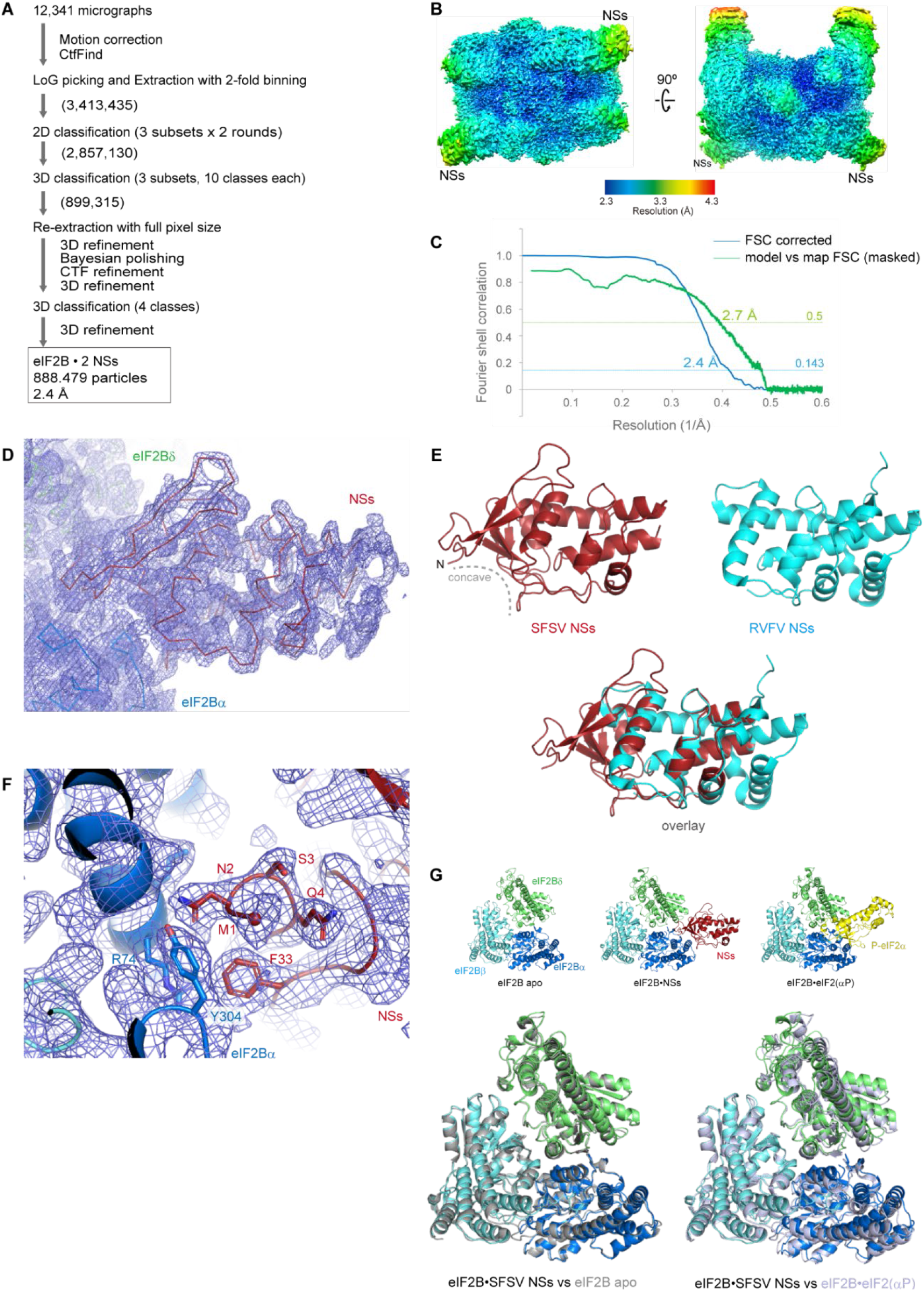
Structural details of the eIF2B•SFSV NSs complex, related to Figure 1 and Table 1. (A) Cryo-EM data processing workflow of image processing. Total particle numbers at each stage are shown in parentheses. (B) Local resolution maps of the eIF2B•SFSV NSs complex from two different views. (C) Fourier shell correlation (FSC) curves of the cryo-EM map. The blue line shows the FSC curve for the 3D reconstruction, and the green line shows the FSC curve calculated between the refined model and the cryo-EM map. (D) EM density map around SFSV NSs. Some unassigned density is also observed in the distal area from the body of eIF2B. (E) Structural comparison of SFSV NSs (maroon) and RVFV NSs (cyan, PDB: 5OOO) (Barski et al., 2017). (F) EM density map around the N-terminus of SFSV NSs. The eIF2Bα subunit accommodates the very N-terminus of SFSV NSs. The sidechain of M1 was not resolved, and therefore its Cα position is shown as a sphere. (G) Comparisons of the conformations of eIF2B regulatory subunits in the eIF2B•SFSV NSs complex, eIF2B apo (PDB: 7D46), and the eIF2B•eIF2(αP) complex structures (PDB: 7D44) (Zyryanova et al., 2021). In the lower panels, the subunits of eIF2B are aligned with their eIF2Bβ subunit C-terminal domains. *Lower left:* Comparison of eIF2B•SFSV NSs (color-coded) and eIF2B apo (grey), *Lower right:* Comparison of eIF2B•SFSV NSs (color-coded) and eIF2B•eIF2(αP) (pale blue).

**Figure S2.**
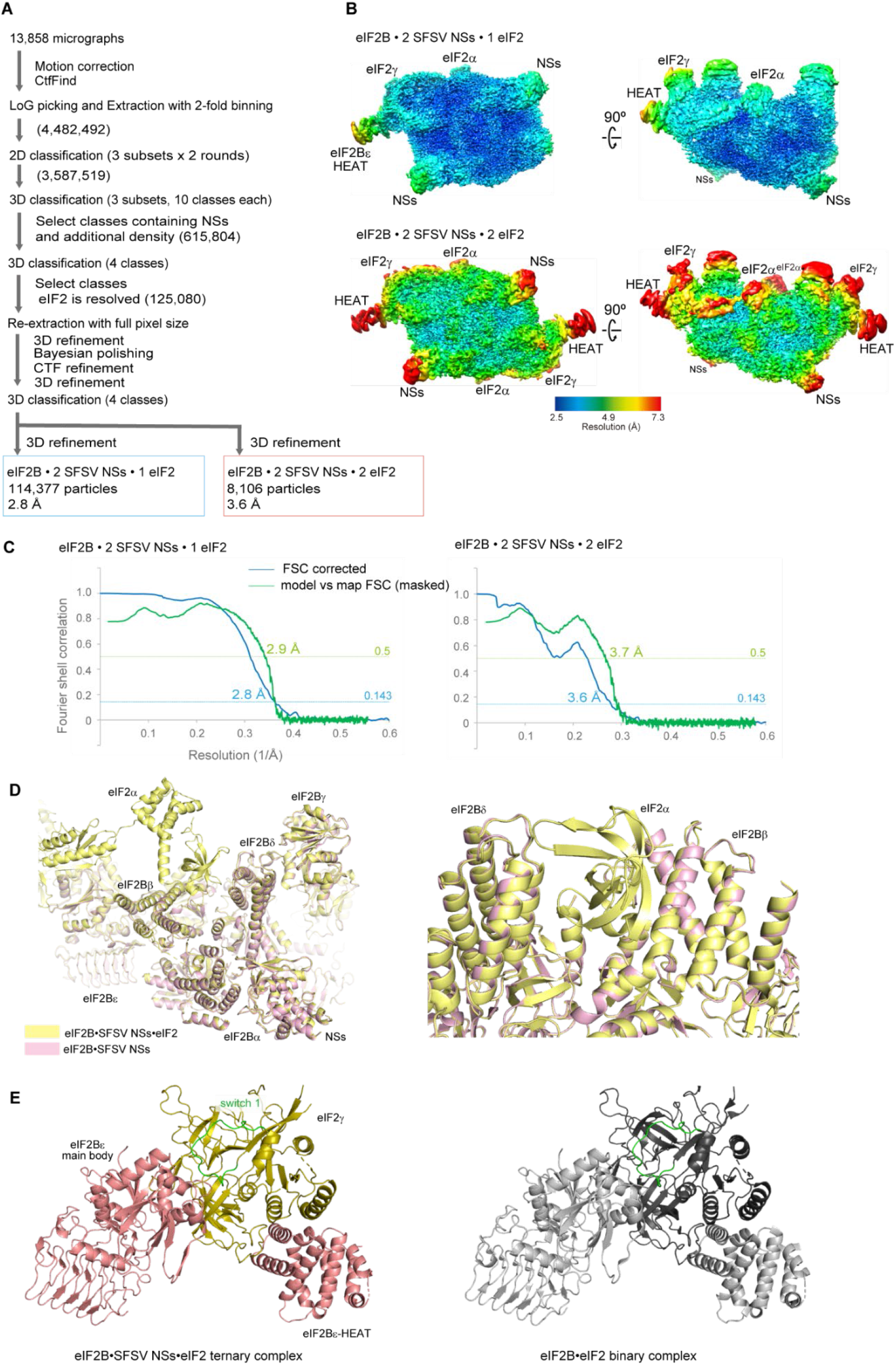
Cryo-EM data processing and structural details of the eIF2B•SFSV NSs•unphosphorylated eIF2 ternary complex, related to Figure 3 and Table 1. (A) Workflow of image processing. Total particle numbers at each stage are shown in parentheses. (B) Local resolution maps of the eIF2B•SFSV NSs•unphosphorylated eIF2 complexes from two different views. (C) Fourier shell correlation (FSC) curves of the cryo-EM maps. The blue lines show the FSC curves for the 3D reconstructions, and the green lines show the FSC curves calculated between the refined models and the cryo-EM maps. (D) Structural comparison of the eIF2B•SFSV NSs complex (pink) and the eIF2B•SFSV NSs•unphosphorylated eIF2 ternary complex (yellow), and close-up view of the interface for the unphosphorylated eIF2 (right panel). Structures were aligned with their eIF2Bβ subunit C-terminal domains. (E) Comparison of the interactions between the eIF2Bε subunit and the eIF2γ subunit in the eIF2B•SFSV NSs•unphosphorylated eIF2 ternary complex (pink and olive) and the eIF2B•eIF2 binary complex structures (grey and black; PDB: 6O85) (Kenner et al., 2019). The switch 1 region of eIF2γ is shown in green.

**Figure S3.**
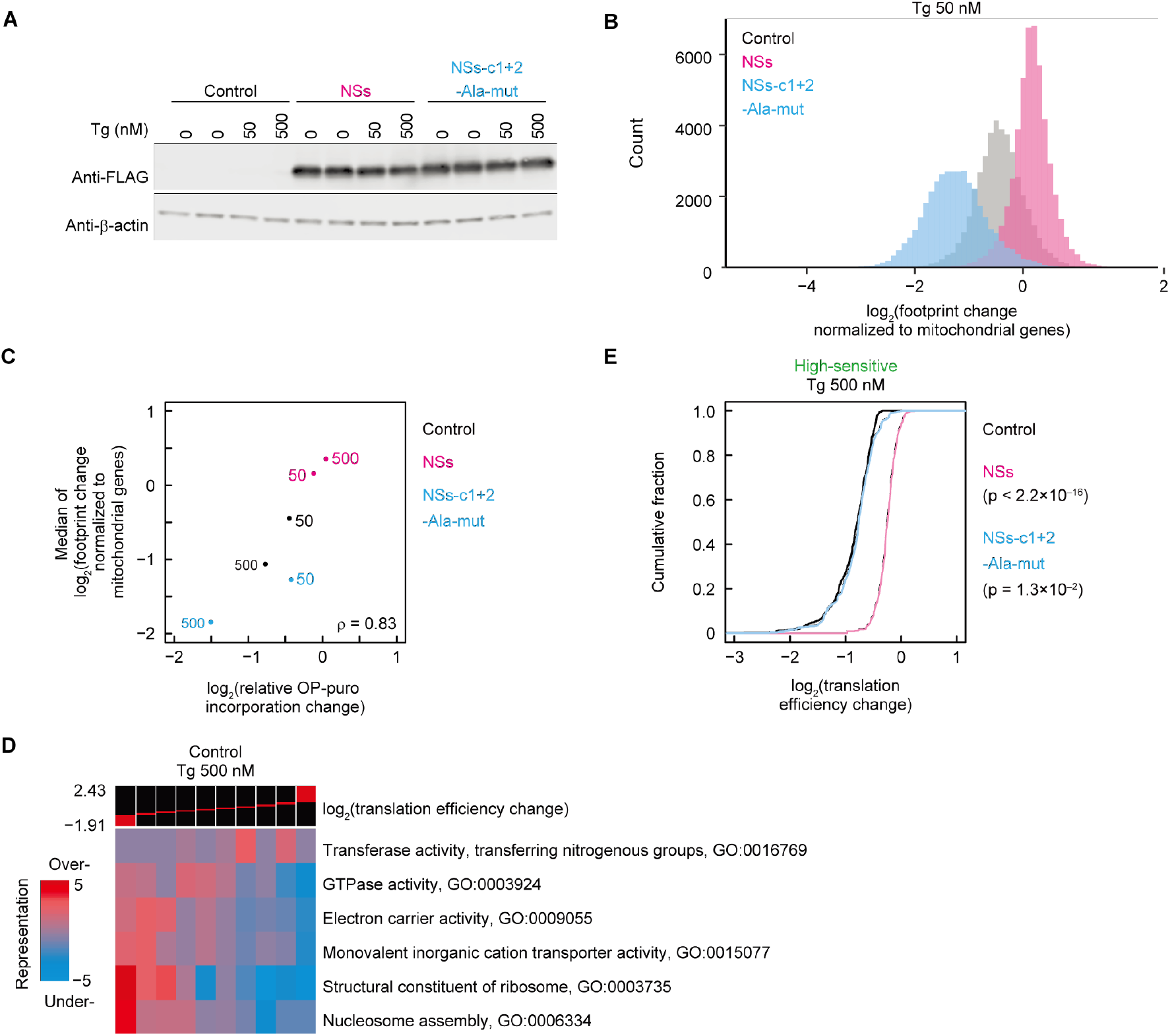
SFSV NSs counteracts the translation repression upon thapsigargin treatment, as revealed by ribosome profiling, related to Figure 4. (A) Western blotting for FLAG-tagged SFSV NSs or SFSV NSs-c1+2-Ala-mut in Tg-treated cells. β-actin was used as a loading control. (B) Histogram of the number of transcripts along the footprint changes in cells treated with 50 nM Tg. Data were normalized to the mean of the footprint changes of mitochondrial genome-encoded genes (used as internal spike-ins). Bin width is 0.1. (C) Comparison between relative OP-puro incorporation change (Figures 4A and 4B) and median of footprint change normalized to mitochondrial ribosome footprints (Figures 4C and S3B). The concentrations of Tg (nM) are shown next to the points. (D) GO analysis of translation efficiency change in control vector-transfected cells treated with 500 nM Tg and visualized by iPAGE (Goodarzi et al., 2009). (E) Cumulative distribution of high-sensitive mRNAs (defined in Figure 4D) along translation efficiency changes in control, SFSV NSs-expressing, or SFSV NSs-c1+2-Ala-mut-expressing cells treated with 500 nM Tg. The p-value was calculated by the Mann-Whitney U test.

**Figure S4.**
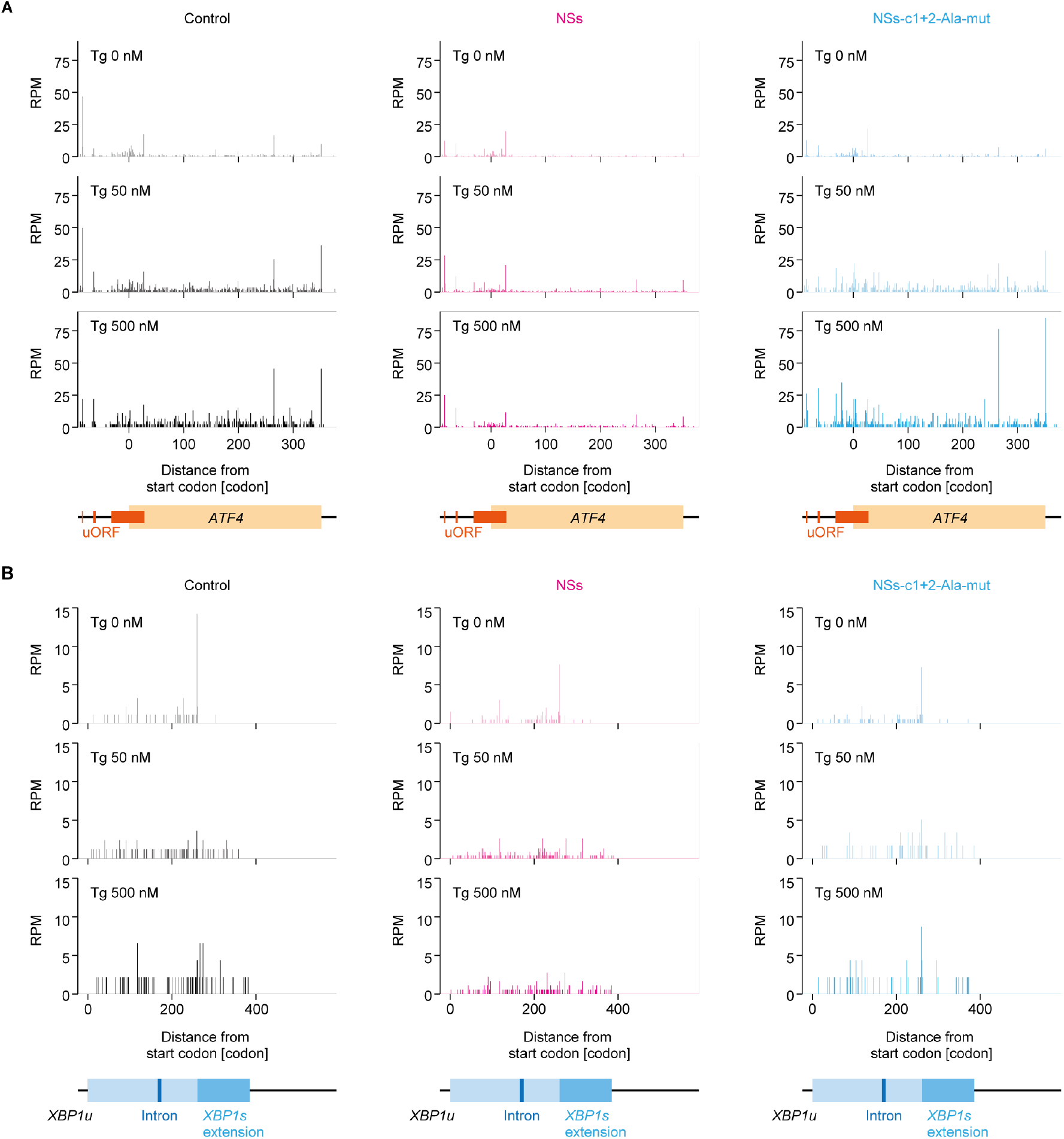
Footprint accumulations on mRNAs activated by thapsigargin, related to Figure 4. The read distributions along *ATF4* (A) and *XBP1u* (B) mRNA in control, SFSV NSs-expressing, or SFSV NSs-c1+2-Ala-mut-expressing cells treated with Tg. The A-site positions of the footprints are shown.

**Table S1.** List of plasmids, related to Figures as indicated.

| Plasmid name | Description | Figures |
| --- | --- | --- |
| pET28-3C-2B1 | For bacterial expression of eIF2Bα, from Kashiwagi et al., 2019 | 1–3, S1, S2 |
| pETDuet-2B4-2B2 | For bacterial expression of eIF2Bδ and $\beta$, from Kashiwagi et al., 2019 | 1–3, S1, S2 |
| pCOLADuet-2B5-2B3 | For bacterial expression of eIF2Bε and $\gamma$, from Kashiwagi et al., 2019 | 1–3, S1, S2 |
| pET28a-d-SFSVNSs | For bacterial expression of SFSV NSs | 1–3, S1, S2 |
| pET28a-d-SFSVNSs-c1-Ala-mut | For bacterial expression of SFSV NSs-c1-Ala-mut (Y5A, Y7A, F33A) | 2B, 2D |
| pET28a-d-SFSVNSs-c2-Ala-mut | For bacterial expression of SFSV NSs-c2-Ala-mut (Y79A, F80A) | 2B, 2D |
| pET28a-d-SFSVNSs-c1+2-Ala-mut | For bacterial expression of SFSV NSs-c1+2-Ala-mut (Y5A, Y7A, F33A, Y79A, F80A) | 2B, 2D |
| pEBMulti-Neo-human-eIF2alpha-PA | For mammalian expression of eIF2α, from Zyryanova et al., 2021 | 2C, 2D |
| pEBMulti-Neo-human-eIF2alpha-S52A-PA | For mammalian expression of eIF2α-S51A, from Zyryanova et al., 2021 | 2C, 2D |
| pEBMulti-Neo-human-eIF2beta | For mammalian expression of eIF2β, from Kashiwagi et al., 2019 | 2C, 2D, 3, S2 |
| pEBMulti-Neo-human-eIF2gamma-FLAG-His8 | For mammalian expression of eIF2γ, from Kashiwagi et al., 2019 | 2C, 2D, 3, S2 |
| pEBMulti-Neo-human-eIF2alpha | For mammalian expression of eIF2α, from Kashiwagi et al., 2019 | 3, S2 |
| pl.18-3xFLAG | For mammalian expression of 3× FLAG peptide | 4, 5, S3, S4 |
| pl.18-SFSVNSs-3xFLAG | For mammalian expression of SFSV NSs protein, from Wuerth et al., 2020 | 4, 5, S3, S4 |
| pl.18-SFSVNSs-c1+2-Ala-mut-3xFLAG | For mammalian expression of SFSV NSs-c1+2-Ala-mut | 4, 5, S3, S4 |
| pCAG-EGFP | For expression of EGFP in rat primary hippocampal neurons & iPS-derived motor neurons | 5 |
| U6 empty plasmid | For expression of EGFP in rat primary hippocampal neurons | 5 |

